# ABCDE approach to victims by lifeguards: how do they manage a critical patient? A cross sectional simulation study

**DOI:** 10.1101/533943

**Authors:** Felipe Fernández-Méndez, Martín Otero-Agra, Cristian Abelairas-Gómez*, Nieves Maria Saez-Gallego, Antonio Rodríguez-Núñez, Roberto Barcala-Furelos

## Abstract

**Introduction:** Decision-making in emergencies is a multifactorial process based on the rescuer, patient, setting and resources. The eye-tracking system is a proven method for assessing decision-making process that has been used in different fields of science. Our aim was to evaluate the lifeguards' capacity to perform the ABCDE (Airway-Breathing-Circulation-Dissability-Exposure) approach when facing a simulated critically ill-drowned victim.

**Methods:** A cross sectional simulation study was designed to assess the skills and sequence of the ABCDE approach by 20 professional lifeguards. They had to assess a victim and act according to his/her clinical status following the ABCDE primary assessment approach. Two kind of variables were recorder: those related to quality of each step of the ABCDE approach; visual behaviour using a portable eye-movement system. The eye-tracking system was the Mobile Eye system (Bedford, USA).

**Results:** None of the study participants was able to complete correctly the ABCDE approach. Lifeguards spent more time in the Circulation step: Airway (15.5±11.1 s), Breathing (25.1±21.1 s), Circulation (44.6±29.5 s), Disability (38.5±0.7 s). Participants spent more time in viewpoints considered as important (65.5±17.4 s) compared with secondary ones (34.6±17.4 s, p = 0.008). This also was represented in the percentage of visual fixations (fixations in important viewpoints: 63.36±15.06; fixation in secondary viewpoints: 36.64±15.06).

**Conclusion:** Professional lifeguards failed to fully perform the ABCDE sequence. Evaluation by experts with the help of eye-tracking technology detected lifeguards' limitations in the assessment and treatment of an eventual critically ill victim. Such deficits should be considered in the design and implementation of lifeguards’ training programmes.

## Introduction

Drowning is a public health problem identified by the WHO as one of the main causes of mortality and morbidity [1], and lifeguards are the professionals with the duty to intervene in an aquatic incident. In addition to environmental water risks, the current occupation and use of aquatic environments for leisure is very popular [2], so other medical complications may also require the attention of lifeguards.

The actions that lifeguards perform when incidents happen in aquatic environments are defined by the drowning timeline, that includes preparation, prevention, rescue and mitigation [3]. Mitigation refers to the skills for the evaluation and treatment of the victim after an incident.

Mitigation of the aquatic incident requires identifying the problem, establishing a preliminary diagnosis and making the appropriate decisions to ease the drowning in a hostile environment [4]. Besides, for each drowned person, it has been estimated that three people receive care in the emergency services [5].

Usually, when talking about drowning, collective thinking associates it with the worst scenario, that is the cardiorespiratory arrest for which the European Guidelines for Resuscitation 2015 recommend to perform cardiopulmonary resuscitation [6]. However, the non-fatal drowning that does not necessarily needs a cardiopulmonary resuscitation, but needs alternative urgent attention is much more prevalent [7]. And it is the one that requires an ABCDE approach to assess the signs and symptoms and to offer an adequate immediate treatment [8].

Unlike other health professionals, for lifeguards 99% of the actions are focused on prevention and rescue and only 1% belong to the care of critically ill patients [7]. However, lifeguards have to be prepared for when this 1% of critical interventions occur, in which’s the decision-making is essential. The decision-making in emergencies is a multifactorial process based on the rescuer, the patient, the setting and the resources, which is difficult to assess since it is an internal process that occurs rapidly [9].

To try to understand this decision-making process, the eye-tracking system was reported as a valid and reliable instrument. It is a proven method that has been used in different fields as sport sciences [10,11] or emergencies [12]. In addition, it is considered a great tool that might positively contribute to improve lifeguards’ skills [13].

Therefore, the aim of the study was to evaluate systematically the decision-making and the capacity in the use of the ABCDE approach by lifeguards.

## Methods

### Sample

A convenience sample of 20 professional lifeguards was invited to participate in this study. Participation was voluntary and authorized through written informed consent. All lifeguards were active professionals trained at the University of Vigo (Spain). Their rescue training was in accordance with regional laws and followed the recommendations for resuscitation of the European Resuscitation Council [14].

### Study design

A cross sectional simulation study was design to evaluate the skills and sequence of the ABCDE approach. This study was approved by the Ethics Committee of the Faculty of Education and Sports Sciences (University of Vigo - Spain) with code 04-1812-17.

### Simulating scenario

The simulating scenario was designed by a multidisciplinary group of experts (emergency doctors, nurses and first-aid coordinators), all of them simulation specialists and instructors. The manikin simulator was programmed according to the values in table 1.

**Table 1.** Simulator programming.

|  |  |
| --- | --- |
| **Airway** | Airway obstruction by tongue fall due to decreased level of consciousness. |
| **Breathing** | Respiratory rate of 40 breaths per minute. Superficial breaths. |
| **Circulation** | Heart rate of 140 beats per minute. Weak radial pulse and strong carotid pulse. Blood pressure of 90/60 mmHg. |
| **Disability** | The simulator did not respond to verbal stimuli. Facing painful stimuli, the patient responded with a scream. |
| **Exposure** | Wound due to erosion in left lower limb with small haemorrhage (distracting wound). |

The participants received the indication that they would enter a room in which they should assess a victim and that they should act according to his/her clinical status following the ABCDE primary assessment approach. The room simulated a beach aid station. The usual material to attend the victim in a real case was available. The simulator remained in the same situation, unchanged, for 10 minutes at which time it went into PCR (see supplementary video). If the participants finished the ABCDE approach before 10 min, the instructor would configure the simulator into cardiac arrest immediately.

The subjective evaluation of the lifeguards' performance was carried out by a BLS instructor and two ALS instructors trained under the ERCGL 2015 guidelines [14,15]. To do this, a checklist was used to evaluate the ABCDE primary assessment based on the recommendations of the Prehospital Trauma Life Support [16].

### Variables

Demographic data such as sex, age, height, weight and body mass index (BMI) were recorded. Afterwards, variables related to the training of the participants were recorded: last training, knowledge of ABCDE approach, have you ever had to perform of ABCDE approach, knowledge of AED and have you done any simulation practice.

Variables regarding to ABCD approach are shown in Table 2. Eye-tracking allowed to collect data of views from the located point were obtained and percentage of time which the located point was viewed from the following areas (defined as viewpoints of great importance): Airway (mouth), Breathing (neck, thorax, abdomen), Circulation (arm, leg, hand, haemorrhage, carotid pulse, radial pulse), Disability (eyes) and Exposure (thermal blanket) and AED.

**Table 2.** Capacity to perform the ABCDE approach.

|  |  |  |  |
| --- | --- | --- | --- |
| **Airway** | Asses airway | Assess the airway correctly | Observe the inside of the mouth for foreign bodies and/or secretion and perform manoeuvre to open the airway or use an oropharyngeal cannula. |
|  |  | Correct order in which the airway is assessed. |  |
| **Breathing** | Assess breathing | Assess the breathing correctly. | Observe if the patient breathes chest symmetry and presence of wounds in the thorax. |
|  |  | Correct order in which the breathing is assessed. |  |
|  |  | Check if the patient is breathing. |  |
|  |  | Check the thoracic symmetry. |  |
| **Circulation** | Assess circulation | Assess the circulation correctly. | Observe the presence of haemorrhage, peripheral and central pulses. |
|  |  | Correct order in which the circulation is assessed. |  |
|  |  | Check central pulse |  |
|  |  | Check peripheral pulse |  |
|  |  | Check the quality of the pulse |  |
|  |  | Check the presence of haemorrhage |  |
| **Disability** | Assess disability | Assess the neurological status correctly. | Evaluate according to the AVPU scale. |
|  |  | Correct order in which the neurological status is assessed. |  |
| **Exposure** | Asses exposure | Assess exposure correctly. |  |
| **Cardiac arrest recognition** |  | Recognise cardiac arrest. |  |
| **Perform CPR with AED** |  | Use AED. |  |

### Instruments

The evaluation was performed on a manikin ALS Simulator (Laerdal, Stavanger, Norway). The gaze fixation variables were obtained through the Mobile Eye gaze tracking system of the ASL laboratories (Bedford, USA). It is based on lightweight glasses that support two cameras: one of them records the scene and the other the point where the vision is focused (extracted by the reflection produced by the cornea and the pupil in a lens). Both signals are registered through its DVCR recording unit and integrated into one via the computer system. This gives us a joint view of the environment observed by the participant and the visual fixations performed. The Mobile Eye system was calibrated using the Eye Vision 2.2.5 software. The resulting videos (supplementary file) have been analysed using the ASL Result Plus Gaze Map software. Both installed in the ACER ASPIRE 5920G laptop (Make INC, Taipei, Taiwan).

## Statistical analysis

The statistical analysis was performed with the Windows statistical package IBM SPSS Statistics version 20 (SPSS, Chicago, Illinois, USA). Continuous variables were described according to measures of central tendency (mean) and dispersion (standard deviation). Categorical variables were described according to absolute and relative frequencies. To verify the normality of the sample, the Kolmogorov-Smirnov test was performed. The Wilcoxon Signed Rank Test was used to verify the differences between the viewpoints of great importance in the ABCDE approach and the unimportant ones, and also to find differences between the percentage of fixations and the percentage of fixation time at each point (significance level of p < 0.05 in both analyses).

## Results

### Characteristics of participants

A total of 18 men and 2 women participated in the study with age of 28.35±6.90 years old, weight of 73.30±8.39 kg, height of 176.40±6.58 cm and BMI of 23.51 ± 2.04 kg•m^−2^.

As for the training of the participants, their last formal training was 31.40±35.40 months before the study. Half of the participants received their last accredited training in a period of 1 year prior to the study. All participants knew what the ABCDE approach was, although 55% had not used it nor had previously performed a simulation scenario test.

### ABCDE approach

None of the study participants managed to complete the primary assessment correctly. Regarding the results of the assessment parts (Fig 1):

**Figure 1.**
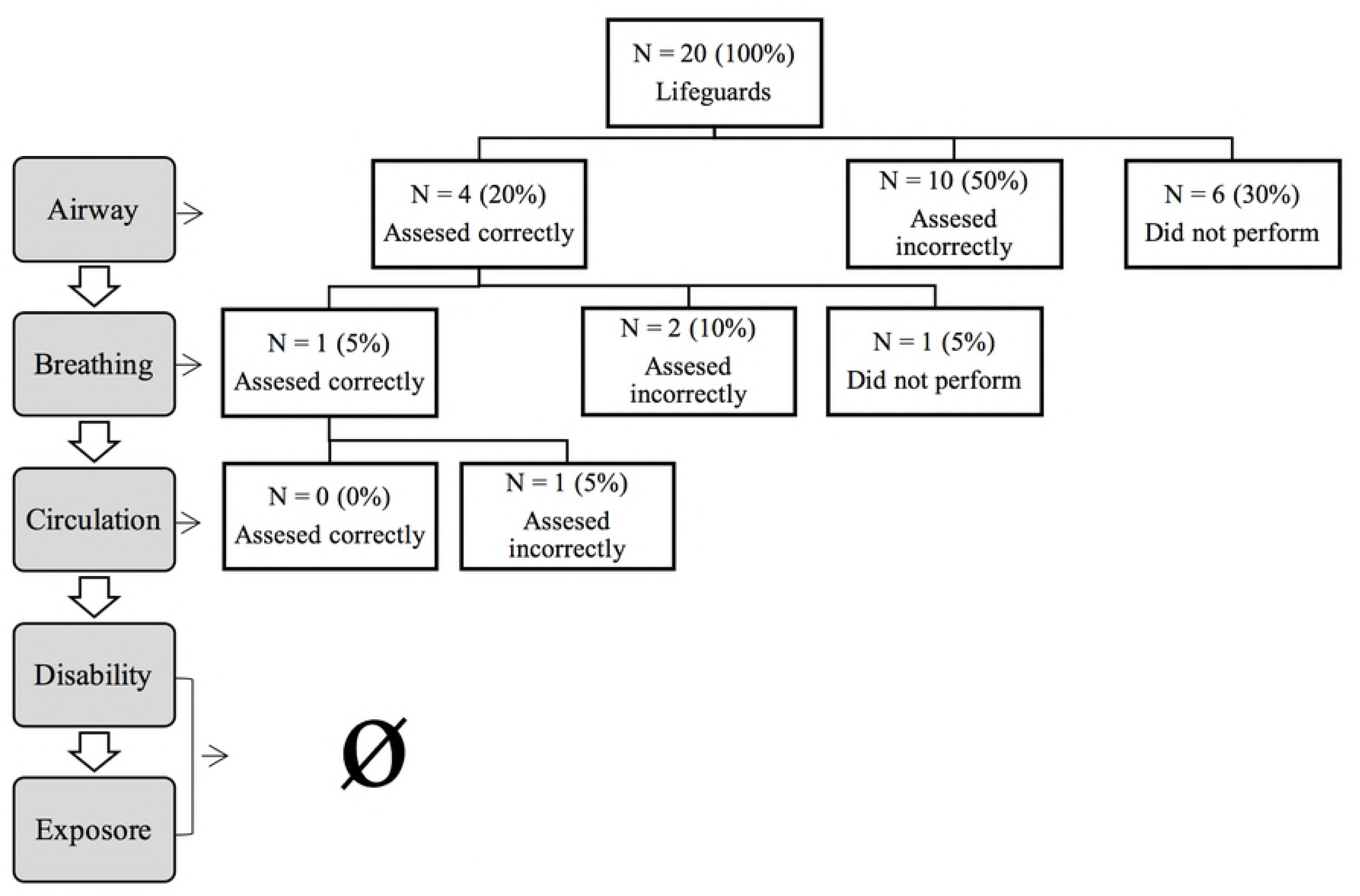
Descriptive analysis about participants’ capacity to perform ABCDE approach.

#### Airway

45% of the participants performed the airway approach as the first step. 30% of the participants did not take this step and half did it incorrectly.

#### Breathing

This part of the primary assessment was carried out by 55% of the participants in second place, and 15% did not contemplate it. 25% of the participants correctly assessed breathing. 80% checked the victim's breathing and 30% checked the symmetries of the victim's thorax.

#### Circulation

60% of the participants assessed the circulation, but none did it correctly and less than half did it in third place (40%). The majority of the participants did not check pulses, and 35% did not perform the assessment of potential haemorrhages.

#### Disability

Only two participants evaluated this step, one in second and another in fourth place.

#### Exposure

25% of the participants performed this step.

#### Cardiac arrest recognition and performance

Half of the participants knew how to recognize the cardiac arrest at the time it took place, but only 35% of the participants used an AED together with CPR.

## Fixations during ABCDE approach (Eye-Tracking)

Data from 4 lifeguards has been lost due to technical issues. No significant differences were observed in any of the vision points when comparing the percentage of fixations and the percentage of fixation time (p > 0.05).

Table 3 and Table 4 show the total percentage of fixings of the most important viewpoints comparing with the unimportant. In addition, the percentage of fixings of each important viewpoint with respect to the total. Regarding to time, time spend in each viewpoint is expressed in percentage and the time spend in each part of the ABCDE approach is shown in seconds.

**Table 3.** Descriptive analysis about fixings performed by participants during ABCDE approach (Total, Airway and Breathing).

| N = 16 (4 lost) |  | Mean (SD) | CI | Wilcoxon Signed Rank Test |
| --- | --- | --- | --- | --- |
| Viewpoints of great importance | Fixings (%) | 63.36 (15.06) | 55.34-71.39 | Total significantly point-views<br>vs.<br>Total insignificantly point-views<br>(p = 0.008) |
|  | Time (%) | 65.45 (17.36) | 56.20-74.70 |  |
| Unimportant viewpoint | Fixings (%) | 36.64 (15.06) | 28.61-44.66 |  |
|  | Time (%) | 34.55 (17.36) | 25.30-43.80 |  |
| Airway |  |  |  |  |
| Mouth | Fixings (%) | 17.58 (9.78) | 12.37-22.79 |  |
|  | Time (%) | 18.16 (11.11) | 12.24-24.08 |  |
| Time in Airway (s) |  | 15.50 (9.17) | 10.20-20.80 |  |
| Breathing |  |  |  |  |
| Neck | Fixings (%) | 4.03 (3.43) | 2.20-5.86 |  |
|  | Time (%) | 4.87 (5.43) | 1.98-7.76 |  |
| Abdomen | Fixings (%) | 5.81 (3.32) | 4.03-7.58 |  |
|  | Time (%) | 7.72 (7.93) | 3.50-11.95 |  |
| Thorax | Fixings (%) | 9.73 (6.98) | 6.01-13.44 |  |
|  | Time (%) | 9.80 (8.47) | 5.29-14.32 |  |
| Time in Breathing (s) |  | 25.12 (21.12) | 14.26-35.98 |  |

Fixings and time of total and each viewpoint in percentage.
Time of each total section of ABCDE approach in seconds.

**Table 4.**
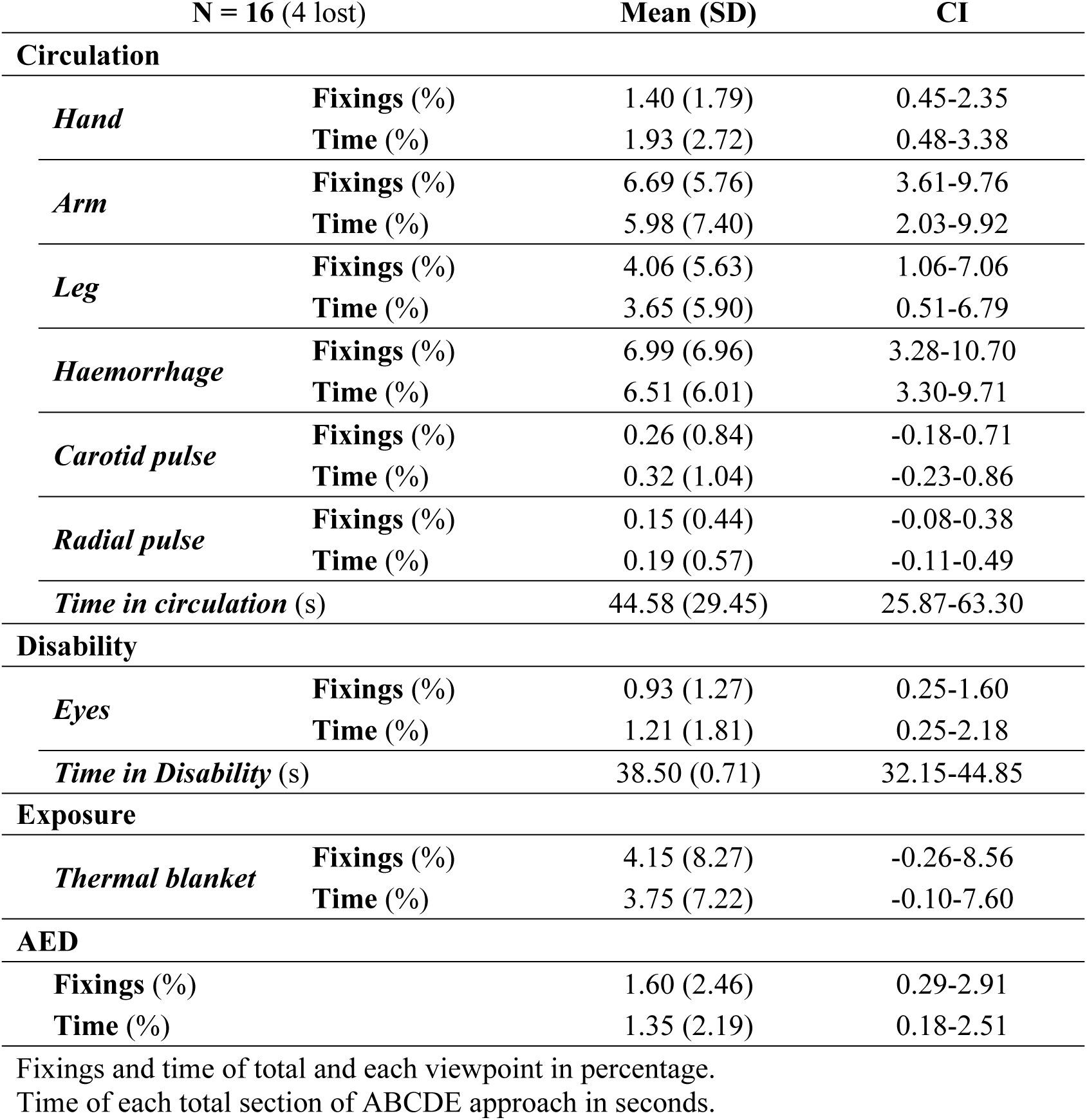
Descriptive analysis about fixings performed by lifeguards during ABCDE approach (Circulation, Disability, Exposure and AED).

| N = 16 (4 lost) |  | Mean (SD) | CI |
| --- | --- | --- | --- |
| **Circulation** |  |  |  |
| *Hand* | **Fixings (%)** | 1.40 (1.79) | 0.45-2.35 |
|  | **Time (%)** | 1.93 (2.72) | 0.48-3.38 |
| *Arm* | **Fixings (%)** | 6.69 (5.76) | 3.61-9.76 |
|  | **Time (%)** | 5.98 (7.40) | 2.03-9.92 |
| *Leg* | **Fixings (%)** | 4.06 (5.63) | 1.06-7.06 |
|  | **Time (%)** | 3.65 (5.90) | 0.51-6.79 |
| *Haemorrhage* | **Fixings (%)** | 6.99 (6.96) | 3.28-10.70 |
|  | **Time (%)** | 6.51 (6.01) | 3.30-9.71 |
| *Carotid pulse* | **Fixings (%)** | 0.26 (0.84) | -0.18-0.71 |
|  | **Time (%)** | 0.32 (1.04) | -0.23-0.86 |
| *Radial pulse* | **Fixings (%)** | 0.15 (0.44) | -0.08-0.38 |
|  | **Time (%)** | 0.19 (0.57) | -0.11-0.49 |
| *Time in circulation (s)* |  | 44.58 (29.45) | 25.87-63.30 |
| **Disability** |  |  |  |
| *Eyes* | **Fixings (%)** | 0.93 (1.27) | 0.25-1.60 |
|  | **Time (%)** | 1.21 (1.81) | 0.25-2.18 |
| *Time in Disability (s)* |  | 38.50 (0.71) | 32.15-44.85 |
| **Exposure** |  |  |  |
| *Thermal blanket* | **Fixings (%)** | 4.15 (8.27) | -0.26-8.56 |
|  | **Time (%)** | 3.75 (7.22) | -0.10-7.60 |
| **AED** |  |  |  |
| **Fixings (%)** |  | 1.60 (2.46) | 0.29-2.91 |
| **Time (%)** |  | 1.35 (2.19) | 0.18-2.51 |

Fixings and time of total and each viewpoint in percentage.
Time of each total section of ABCDE approach in seconds.

Around 60% of the fixations and fixation time were devoted to important areas of vision for the ABCDE approach (Fig 2), with significant differences being found when compared with unimportant vision areas (p < 0.008).

**Figure 2.**
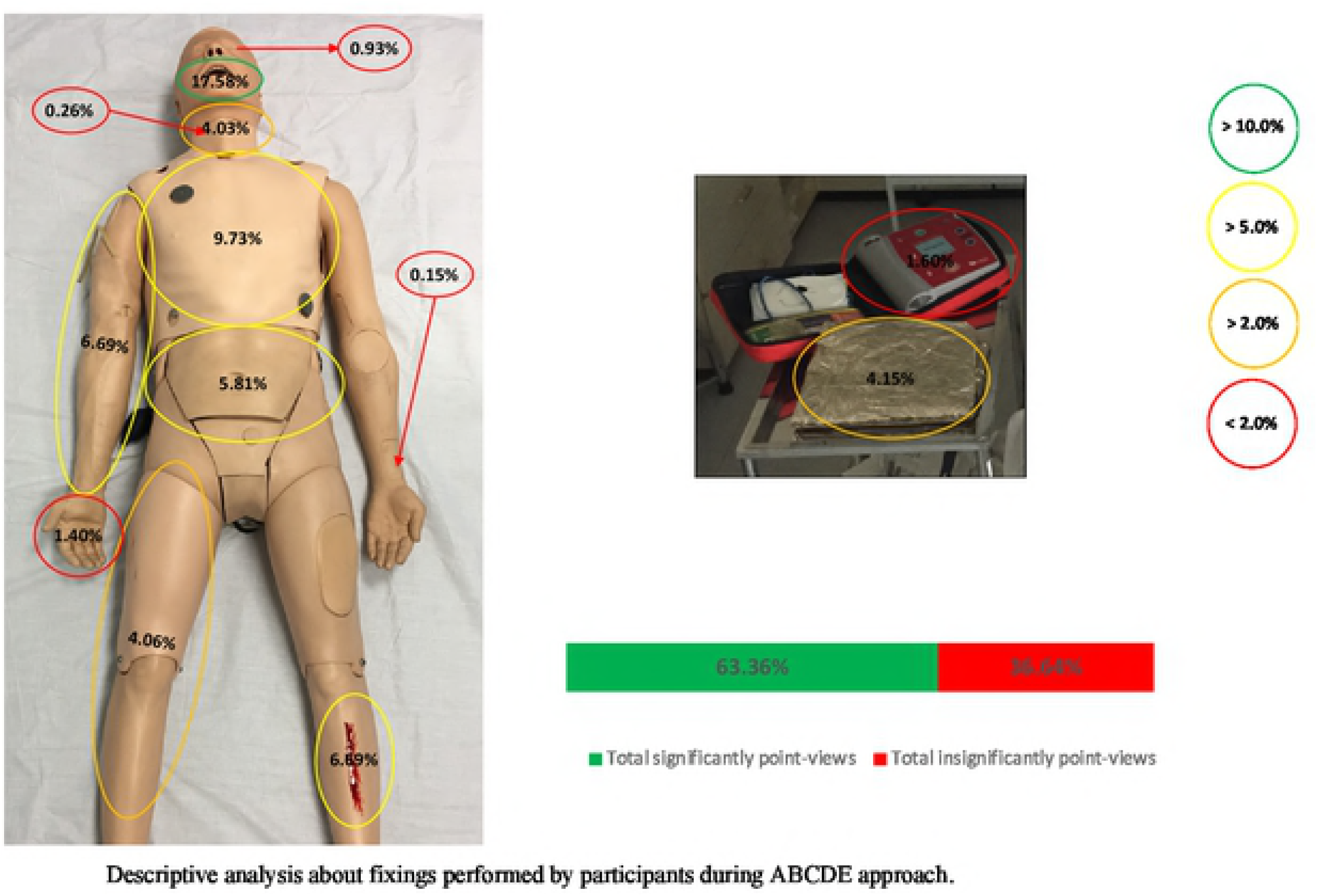
Descriptive analysis about fixings performed by participants during ABCDE approach.

The "Circulation" approach was the one that required more time to carry out its assessment (44.6 s), followed by "Disability" (38.5 s) and "Breathing" (25.1 s). The steps that required less time were "Exposure" (3.8 s) and "Airway" (15.5±11.1 s).

### Airway

A 17.58% of fixations was required to assess the mouth-airway.

### Breathing

The assessment of the thorax was the stage that required the highest percentage of fixations on the total number of fixations (9.73%). The assessment of the neck was the one with the lowest percentages (4.03%). The assessment of the abdomen shows a fixation percentage of 5.81%.

### Circulation

The assessment of bleeding was the one that required the highest fixation percentages (6.99%), followed by the arm (6.69%). The assessment of the legs obtained percentages of 4.06%, while the evaluation of the hand, carotid pulse and radial pulse obtained percentages lower than 2%.

### Disability

The eye assessment obtained fixation percentages of 0.93%.

### Exposure

The thermal blanket obtained higher percentages (4.15%).

## Discussion

Simulation training is a well-recognised educational tool in medical and emergency personnel education that allow improvement of technical and non-technical skills [17–19]. Eye-tracking technology was reported as effective instrument in order to complement the simulation training and the subjects performance assessment [20,21]. Recent publications show its effectiveness assessing or evaluating in simulating scenarios related to anaphylaxis or paediatric trauma [12,22]. In fact, the accuracy of this tool also resulted in its using in real situations [23].

In our study, eye-tracking was used to analyse the lifeguards’ ABCDE approach skills. Professional lifeguards showed differences when it comes to recognizing and treating a critically ill patient in a simulated scenario. Although all the participants had theoretical training and knew what was and how was applied the primary assessment, they had not received simulation training and half of them had not used the primary assessment in a real situation. Although there is no clear evidence that training "life support in trauma" has an impact on the results of trauma victims, there is evidence that educational initiatives improve knowledge about what to do in emergency situations [24].

Most of rescuers of this study evaluated the Airway (14 of 20: 70%) and Breathing (17 of 20: 85%) although only 4 (20%) and 5 (25%) assessed both correctly. Only 40% evaluated Circulation. Of the subjects who performed the evaluation of Circulation, only 3 took a central pulse and 2 checked a peripheral pulse. The assessment of Circulation was the step that required the longest time assessment (45 s). Maybe this was due to the haemorrhage that the simulator presented in the lower limb as a distracting factor introduced by the researchers. The haemorrhage was small and produced by an erosion. The majority of the participants did not consider assessing pulses neither peripheral nor central. This could be due to that the Guidelines of the European Resuscitation Council as well as the Guidelines of the American Heart Association of basic life support insist that taking the pulse is not a necessary measure to establish the diagnosis of cardiorespiratory arrest [14,25]. However, the simulator did not present a PCR and required a Circulation evaluation.

The application of a structured assessment system has become the norm in trauma. This approach to the early recognition and treatment of life-threatening injuries has been trained in trauma courses for decades [26,27]. In the study by Olgers et al. [28] in which they investigated the use of the ABCDE approach by emergency doctors they observed that this approach was used in 26% of patients. When ABCDE approach was used it was done with high scores (83%). The reason why the doctors decided not to use this assessment approach was because of the general clinical impression, the vital signs registered by nursing or that the reason for the consultation does not suggest an unstable patient [28]. In another study conducted in a hospital emergency department, it was found that only 52% of patients were evaluated with the ABCDE approach, and 17% were fully evaluated with precision [29].

In view of our results, it seems that the lifeguards are not competent performing the ABCDE approach in a simulated scenario. With respect to the visual fixations, the lifeguards were able to maintain an adequate attention and fixation, which means that they are focused on looking. Around 60% of the fixations and the time of visual fixations were dedicated to important areas of vision for the ABCDE approach. This is independent of the decisions made at each moment because the eyes could be focusing on the important parts and not knowing the decisions to make.

## Practical implications

The poor results found in this study are relevant and useful to re-design and to re-organize the training of lifeguards in two ways. For one hand, more efforts are needed to train in terms of ABCDE approach considering the correct order of the different steps and its relevance in the sequence. On the other hand, ABCDE approach comprises a set of different skills and knowledges whose competency might decrease over time. Hence, lifeguards’ training should consider this aspect and periodically refreshing re-training should be mandatory to maintain the skills quality.

## Limitations

This is a simulation study; therefore, a real intervention could generate different results, which we suppose it might be even worse than the observed in our trial. The lack of experience of the participants in the use of the simulation methodology could be a limiting factor. The subjects sample was small, so the results should be taken with caution. Also, the duration of the scenario could be a limiting factor as well because we do not know what could have happened in the case that the scenario had lasted longer. The participants knew that they were under observation and this may have modified their performance.

## Conclusions

Professional lifeguards showed skill limitations in a simulated scenario that required ABCDE approach. Evaluation by experts and eye-tracking technology detected many gaps in the assessment and treatment of an eventual critically casualty. Such deficits should be considered in the re-design and implementation of lifeguards’ training programmes.

